# WSL5, a pentatricopeptide repeat protein, is essential for chloroplast biogenesis in rice under cold stress

**DOI:** 10.1101/239905

**Authors:** X. Liu, J. Lan, Y.S. Huang, P.H. Cao, C.L. Zhou, Y.K. Ren, N.Q. He, S.J. Liu, Y.L. Tian, T.L. Nguyen, L. Jiang, J.M. Wan

## Abstract

Chloroplasts play an essential role in plant growth and development, and cold has a great effect on chloroplast development. Although many genes or regulators involved in chloroplast biogenesis and development have been isolated and characterized, identification of novel components associated with cold is still lacking. In this study, we reported the functional characterization of *white stripe leaf 5* (*wsl5*) mutant in rice. The mutant developed white-striped leaves during early leaf development and was albinic when planted under cold stress. Genetic and molecular analysis revealed that WSL5 encodes a novel chloroplast-targeted pentatricopeptide repeat protein. RNA-seq analysis showed that expression of nuclear-encoded photosynthetic genes in the mutant was significantly repressed, and expression of many chloroplast-encoded genes was also significantly changed. Notably, the *WSL5* mutation caused defects in editing of *rpl2* and *atpA*, and in splicing of *rpl2* and *rps12*. Chloroplast ribosome biogenesis was impaired under cold stress. We propose that *WSL5* is required for normal chloroplast development in rice under cold stress.

## Introduction

Cold is an important environmental factor affecting chloroplast development and growth in juvenile plants and sudden low-temperature periods that often occur during early seedling development in spring can directly affect production (Kusumi and Iba, 2014). Rice seedlings are susceptible to cold stress with an impact that ultimately affects grain yield (Liu *et al*. 2013). Therefore, cold stress is a common problem that affects grain production, and rice varieties with increased cold tolerance are preferred (Zhao *et al*. 2017). Many studies have suggested that plants can regulate early chloroplast development under cold stress. In certain virescent mutants the chlorophyll content in young leaves is low, but gradually increases to normal levels as they develop (Yoo *et al*. 2009). Temperature-sensitive virescent mutants were used to study mechanisms regulating chloroplast development in seedlings under cold stress conditions, and many genes were identified, such as *V3, St1*, *OsV4, TCD9* and *TSV* (Yoo *et al*. 2009; Gong *et al*. 2014; Jiang *et al*. 2014; Sun *et al*. 2017). However, the mechanisms of chloroplast development in rice seedlings under cold stress remain poorly understood.

Chloroplasts are essential for photosynthesis and have crucial roles in plant development and growth by fixation of CO_2_ and biosynthesis of carbon skeletons as well as other physiological processes (Jarvis and López-Juez, 2013). Formation of a photosynthetically active chloroplast from a proplastid is controlled by both nucleus-encoded polymerase (NEP) and plastid-encoded polymerase (PEP) and is accompanied by rapid development of the thylakoid membrane (Yu *et al*. 2014). NEP is a simple protein that is responsible for transcription of genes encoding plastidic PEP subunits, ribosomal proteins, and other plastidic “housekeeping” proteins (Liere *et al*. 2011). PEP, on the other hand, is a large, complex protein with many transiently attached peripheral subunits that participate in photosynthesis at the later stages of chloroplast development (Yu *et al*. 2014).

Chloroplast RNAs need to be processed to become functional rRNAs and mRNAs. Many of the processing factors for RNA cleavage, splicing, editing and stability are RNA-binding proteins (Tillich and Krause, 2010). All are coded by the nuclear genome. One family of RNA-binding proteins has pentatricopeptide repeats (PPR) and usually carries out specific RNA processing in chloroplasts, a feature first recognized from the *Arabidopsis thaliana* genome sequence (Stern *et al*. 2010; Shikanai and Fujii, 2013). PPR proteins are defined by a tandem array of a PPR motif consisting of 35 amino acids. In higher plants, the PPR family contains many members, with 450 in Arabidopsis and 655 in rice (O’Toole *et al*. 2008). The functions of PPR proteins are well characterized (Stern *et al*. 2010; Shikanai and Fujii, 2013). Chloroplast-targeted PPR proteins were characterized as being involved in regulating RNA splicing, RNA editing, RNA cleavage, RNA stability, and RNA translation during plant development and growth (Yu *et al*. 2009; Ichinose *et al*. 2012). Several PPR genes in rice, such as *YSA*, *OsV4, WSL, ALS3, OspTAC2*, and *WSL4*, were reported to function in chloroplast biogenesis, RNA editing, RNA splicing and chloroplast development (Su *et al*. 2012; Gong *et al*. 2014; Tan *et al*. 2014; Lin *et al*. 2015; Wang *et al*. 2016; Wang *et al*. 2017). A PPR mutant in rice, *ysa*, develops albinic leaves before the three leaf stage, but the plants gradually turn green and recover to normal green at the six leaf stage (Su *et al*. 2012). *WSL* encodes a rice PPR protein that targets the chloroplasts and plays an essential role in splicing the chloroplast transcript *rpl2* (Tan *et al*. 2014). The P-family PPR mutant *wsl4*, which exhibits white-striped leaves before the 5-leaf stage, has defective chloroplast RNA group II intron splicing (Wang *et al*. 2017). However, the functions, substrates and regulatory mechanisms of many PPR proteins in rice remain to be elucidated.

In this study, we isolated and characterized a rice mutant *wsl5* that develops white-striped leaves at the early seedling stage; *wsl5* is albinic at low temperatures. *WSL5* encodes a P-family PPR protein containing an RNA recognition motif at its N terminus and 15 PPR motifs at its C terminus. WSL5 locates to chloroplasts and is essential for chloroplast ribosome biogenesis under cold stress. We showed that RNA editing sites of *rpl2* and *atpA* were not edited in the mutant and plastid-encoded genes *rpl2* and *rps12* were not efficiently spliced in the *wsl5* mutant. Our results provide insight into the function *WSL5* in rice chloroplast development under cold stress.

## Materials and methods

### Plant materials and growth conditions

The *wsl5* mutant was selected from an ethyl methane sulfonate (EMS) mutagenesis mutant pool of the subspecies *indica* cultivar Nanjing 11. Seeds of the WT and *wsl5* plants were grown in a growth chamber under 16 h of light/8 h of darkness at constant temperatures of 20, 25, and 30°C. The third leaves at about 10-days post planting were used for nearly all analyses. To map the *WSL5* locus, we constructed an F_2_ population derived from a cross of the *wsl5* mutant and Dongjin(*japonica*).

### Pigment determination and transmission electron microscopy

Wild-type and *wsl5* mutant seedlings were grown in the field. Fresh leaves were collected and used to determine chlorophyll contents using a spectrophotometer according to the method of Arnon (1949). Briefly, 0.2 g of leaf tissue were collected and marinated in 5 ml of 95% ethanol for 48 h in darkness. The supernatants were collected by centrifugation and were analysed with a DU 800 UV/Vis Spectrophotometer (Beckman Coulter) at 665, 649 and 470 nm, respectively.

Transmission electron microscopy was performed according to the method of Wang *et al*. (2016). Briefly, fresh leaves were collected and cut into small pieces, fixed in 2.5% glutaraldehyde in a phosphate buffer at 4°C for 4 h, further fixed in 1% OsO_4_, stained with uranyl acetate, dehydrated in an ethanol series, and finally embedded in Spurr’s medium prior to ultrathin sectioning. The samples were observed using a Hitachi H-7650 transmission electron microscope.

### Map-based cloning and complementation of WSL5

Genetic analysis was performed using an F_2_ population (*wsl5*/Nanjing11); 654 plants with the recessive mutant phenotype were used for genetic mapping. New SSR/Indel markers were developed based on the sequences of Nipponbare and 93–11(*indica*) genomes (http://www.gramene.org/). The *WSL5* locus was narrowed to a 180 kb region flanked by InDel markers Y18 and Y47 on the long arm of chromosome 4 (Table S2).

For complementation of the *wsl5* mutation, a 2,706 bp WT CDS fragment and an ~2 kb upstream sequence were amplified from variety Nanjing 11. They were cloned into the binary vector pCAMBIA1390 to generate the vector pCAMBIA1390-*WSL5*. This vector was introduced into *Agrobacterium tumefaciens* strain EHA105, which was then used to infect *wsl5* mutant calli.

### Sequence analysis

Gene prediction and structure analysis were performed using the GRAMENE database (www.gramene.org/). Homologous sequences of WSL5 were identified using the Blastp search program of the National Center for Biotechnology Information (NCBI, www.ncbi.nlm.nih.gov/). Multiple sequence alignments were conducted with DNAMAN.

### Subcellular localization of WSL5 protein

For subcellular localization of WSL5 protein in rice protoplasts, the coding sequence of WSL5 was amplified and inserted into the pAN580 vector. The cDNA fragments were PCR-amplified using primer pairs shown in Table S2. Protoplasts were isolated from 10-day-old 9311 seedlings. Transient expression constructs were separately transformed into rice protoplasts and incubated in the darkness at 28°C for 16 h before examination (Chen *et al*. 2006). GFP fluorescence was observed using a confocal laser scanning microscope (Zeiss LSM 780).

### Quantitative RT-PCR analysis

Total RNA was isolated using the RNA prep pure plant kit (TIANGEN, Beijing). First-strand cDNA was synthesized using random hexamer primers (TaKaRa) for chloroplast-encoded genes and oligo(dT)_18_ (TaKaRa) for nuclear encoded genes, and reverse transcribed using Prime scriptase (TaKaRa). Real-time PCR (RT-PCR) was performed using an ABI 7500 real-time PCR system with SYBR Green MIX and three biological repeats. Primers used for RT-PCR are listed in Table S2. The rice *Ubiquitin* gene was used as an internal control.

### RNA analysis

Total RNA was isolated from 10-d-old seedlings of wild type and *wsl5* grown in C30 and C20 conditions using the RNA prep pure plant kit grown in the field. RNA samples were diluted to 10 ng/mL and analyzed by an Agilent 2100 bioanalyzer. The RNA 6000 Nano Total RNA Analysis Kit (Agilent) was used for analysis.

### RNA editing sites and RNA splicing analysis

Specific cDNA fragments were generated by RT-PCR amplification following established protocols (Takenaka & Brennicke 2007). The cDNA sequences were compared to identify C to T changes resulting from RNA editing. For RNA splicing analysis, the chloroplast gene with at least one intron was selected and amplified using RT-PCR with primers flanking the introns. The primers used for RNA editing and splicing analysis were obtained as reported previously (Tan *et al*. 2014; Zhang *et al*. 2017).

### Protein extraction, SDS-PAGE, and western blotting

Leaf material was homogenized in lysis buffer (25 mM Tris–HCl, pH 7.6, 0.15 M NaCl, and 2% sodium dodecyl sulfate (SDS), 0.01% 2-mercaptoethanol). Sample amounts were standardized by fresh weight. The protein samples were separated by 10% SDS–polyacrylamide gel electrophoresis (PAGE). After electrophoresis, the proteins were transferred onto PVDF membranes (Millipore) and incubated with specific antibodies. Signals were detected using an ECL Plus Western Blotting Detection Kit (Thermo) and visualized by an imaging system (ChemiDocTMX-RS; Bio-Rad).

### Yeast two-hybrid analysis

The coding sequences of five rice MORFs were amplified with primers listed in Zhang *et al*. (2017), and then MORFs and WSL5 were cloned into the pGAD-T7 or pGBK-T7 vectors, respectively. Yeast two-hybrid analysis was performed using the Clontech (Clontech, www.clontech.com) two-hybrid system, following the manufacturer’s instructions.

### RNA-seq analysis

Total RNA was extracted from 10-day-old wild type and *wsl5* seedlings grown at different temperatures. mRNA was enriched from total RNA using oligo-(dT) primer and Ribo-Zero rRNA Removal Kits for chloroplast-encoded genes. cDNA was synthesized using random hexamer primers. The library was constructed and sequenced using an Illumina Hisequation 2000 (TGS, Shenzhen). Totals of 45 million reads of genes from wild type and 42 million from *wsl5* were obtained. The significance of differentially expressed genes (DEGs) were determined by using |log_2_ (Fold Change)| >1 and q values<0.05. Gene Ontology (http://www.geneontology.org/) analyses were performed referring GOseq (Young *et al*. 2010). Pathway enrichment analysis was performed using the Kyoto Encyclopedia of Genes and Genomes database (Kanehisa *et al*. 2008).

## Results

### Characterization of the wsl5 mutant

To identify genetic factors regulating chloroplast development in rice, we used the *wsl5* mutant obtained in an EMS mutant pool of Nanjing 11(*indica*). Seedlings of *wsl5* exhibited a white-striped leaf phenotype up to the four-leaf stage under field conditions (Fig. 1A, B). Normal green leaf development occurred thereafter. Chlorophyll and carotenoid contents in leaves of *wsl5* mutant were much lower than in wild type (WT) before the five-leaf stage, but were subsequently similar to the WT (Supplementary Fig. S1). Major agronomic traits of the *wsl5* mutant at maturity, such as plant height and seed size, were indistinguishable from those of WT plants (Fig. 1C and Supplementary Table S1). Chlorophyll (Chl a, Chl b) and carotenoid contents were reduced in *wsl5* mutant seedlings (Fig. 1D). To examine whether lack of photosynthetic structure was accompanied by ultrastructural changes in chloroplasts of *wsl5* mutant, we compared the ultrastructure of chloroplasts in white and green sectors of *wsl5* mutant leaves and normal WT leaves by transmission electron microscopy (TEM). Cells in WT leaves and green sectors in leaves of *wsl5* had normal chloroplasts displaying structured thylakoid membranes composed of grana connected by stroma lamellae (Fig. 1E, H). However, the chloroplasts within the white sectors in the *wsl5* mutant were abnormal (Fig. 1I, J). The results suggested that the *WSL5* had a role in chloroplast development in juvenile plants.

**Fig. 1.**
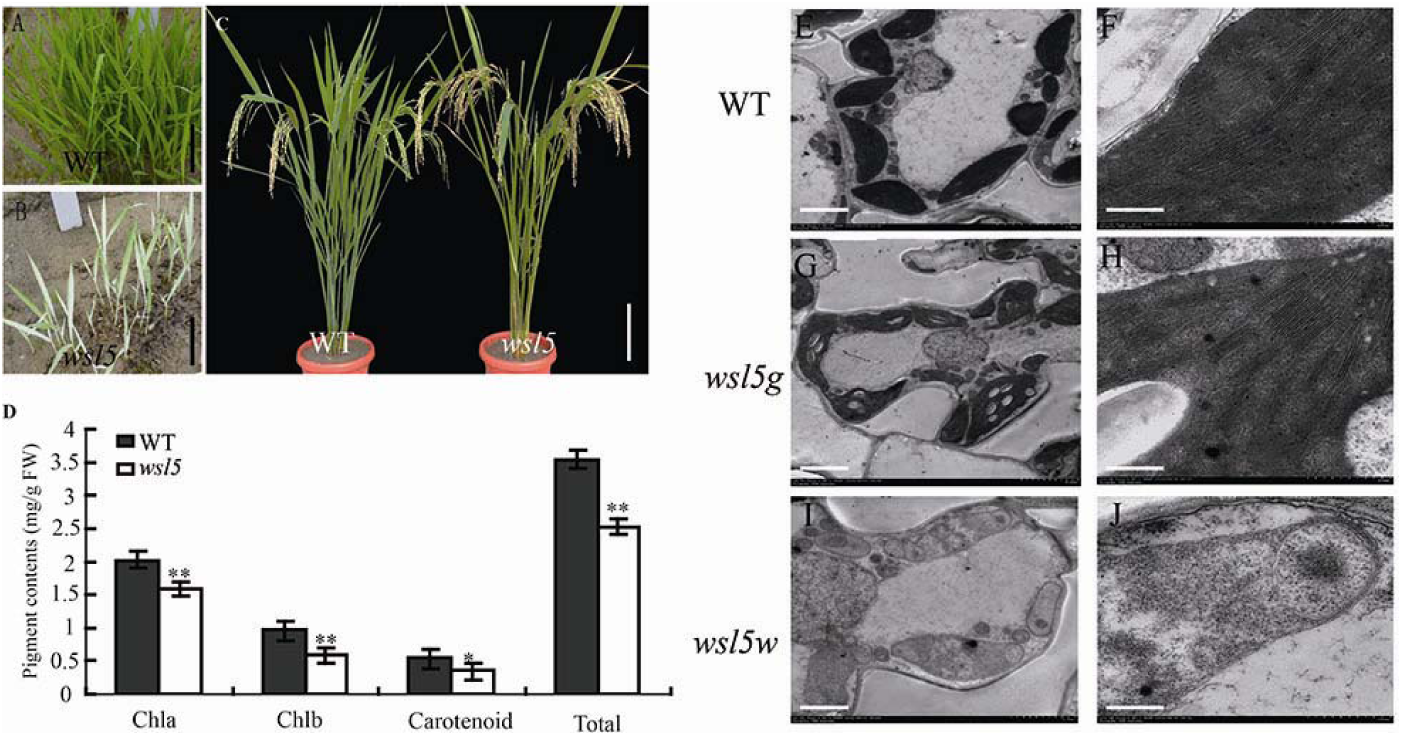
Phenotypic characteristics of *wsl5* mutant. (A-B) Phenotypes of WT and *wsl5* mutant seedlings in the field 20 days after seeding. (C) Phenotypes of WT (left) and *wsl5* (right) plants at maturity. (D) Leaf pigment contents of field-grown WT and *wsl5* seedlings at 20 days after seeding. (E-F) Mesophyll cells in wild-type plants showing normal, well ordered chloroplasts. (G-H) Chloroplasts from green sectors of *wsl5* seedlings were indistinguishable from those of WT. (I-J) Cells from white sectors of the mutants displayed abnormalities, including vacuolated plastids and lack of organized thylakoid membranes. Scale bar: 1 cm in (A-B), 10 cm in (C), 1 μm in (E, G, I), 500 nm in (F, H, J). (Student’s t-test, \*\**P* < 0.01, * *P* < 0.05).

### The wsl5 phenotype was temperature-sensitive

To verify whether the *wsl5* mutant was affected by temperature, *wsl5* and WT seedlings were produced in growth chambers under constant temperatures of 20°C, 25°C, 30°C (C20, C25, C30). Leaves of the *wsl5* mutant were albinic at 20°C (Fig. 2E) and the plants died. Chlorophyll (Chl) was not detectable in the leaves (Fig. 2F). At 25°C *wsl5* mutant developed leaves with white-stripes and chlorophyll was present at reduced levels relative to WT (Fig. 2C, D). At 30°C the mutant exhibited almost the same phenotype as the WT (Fig. 2A) and contained similar Chl contents (Fig. 2B). These results indicated that *wsl5* was sensitive to low temperatures and that *WSL5* protected chloroplast development from cold stress.

**Fig. 2.**
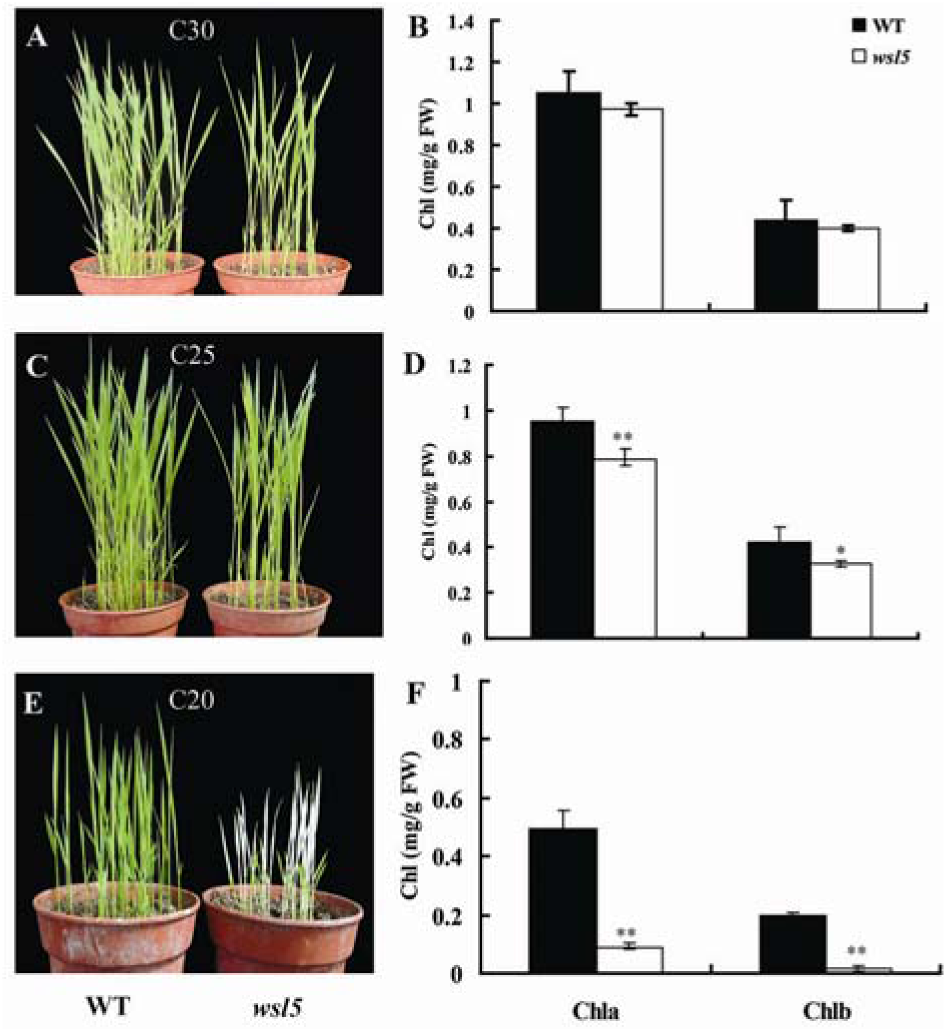
The *wsl5* phenotype is temperature-sensitive. (A, C, E) Phenotypes of WT and *wsl5* seedlings produced at different constant temperatures. C20, C25, and C30, refer to 20, 25, and 30°C respectively. (B, D, F) Leaf pigment contents of WT and *wsl5* seedlings grown at different temperatures. (Student’s t-test, \**P* < 0.05, \*\**P* < 0.01).

We also examined the ultrastructure of the chloroplasts in mesophyll cells of the WT and *wsl5* plants. At 30°C all WT and *wsl5* plants displayed normal chloroplasts with well-developed lamellar structures and with normally stacked grana and thylakoid membranes (Supplementary Fig. S2A-D). At 20°C, WT developed large starch grains and chloroplasts with normal thylakoids (Supplementary Fig.S2E, F), whereas leaf cells from albinic sectors in *wsl5* had no chloroplasts (Supplementary Fig. S2G, H). The results suggested that *WSL5* protected chloroplasts from damage caused by cold stress in wild-type seedlings.

### Map-based cloning of the WSL5 allele

Genetic analysis showed that the white stripe phenotype in the *wsl5* mutants was controlled by a single recessive nuclear locus. To identify the location of the *WSL5* locus, 20 F_2_ individuals with the mutant phenotype derived from a cross between *wsl5* and Dongjin (*japonica*) were used. The *WSL5* locus was located to a 2.65 Mb region flanked by simple sequence repeat (SSR) markers RM8217 and RM559 on the long arm of chromosome 4. It was further delimited to a 180 kb region between indels Y17 and Y47 using 654 F_2_ plants with mutant phenotype. Twenty-two open reading frames (ORFs) were predicted in the region from published data (http://www.gramene.org/; Fig. 3A). Sequence analysis of the region showed that only one ORF encoding a pentatricopeptide repeat protein differed between WT and *wsl5* (Fig. 3B). A SNP (T to C) located in the conserved region caused a leucine to proline amino acid substitution in the mutant (Fig. 3B-C).

**Fig. 3.**
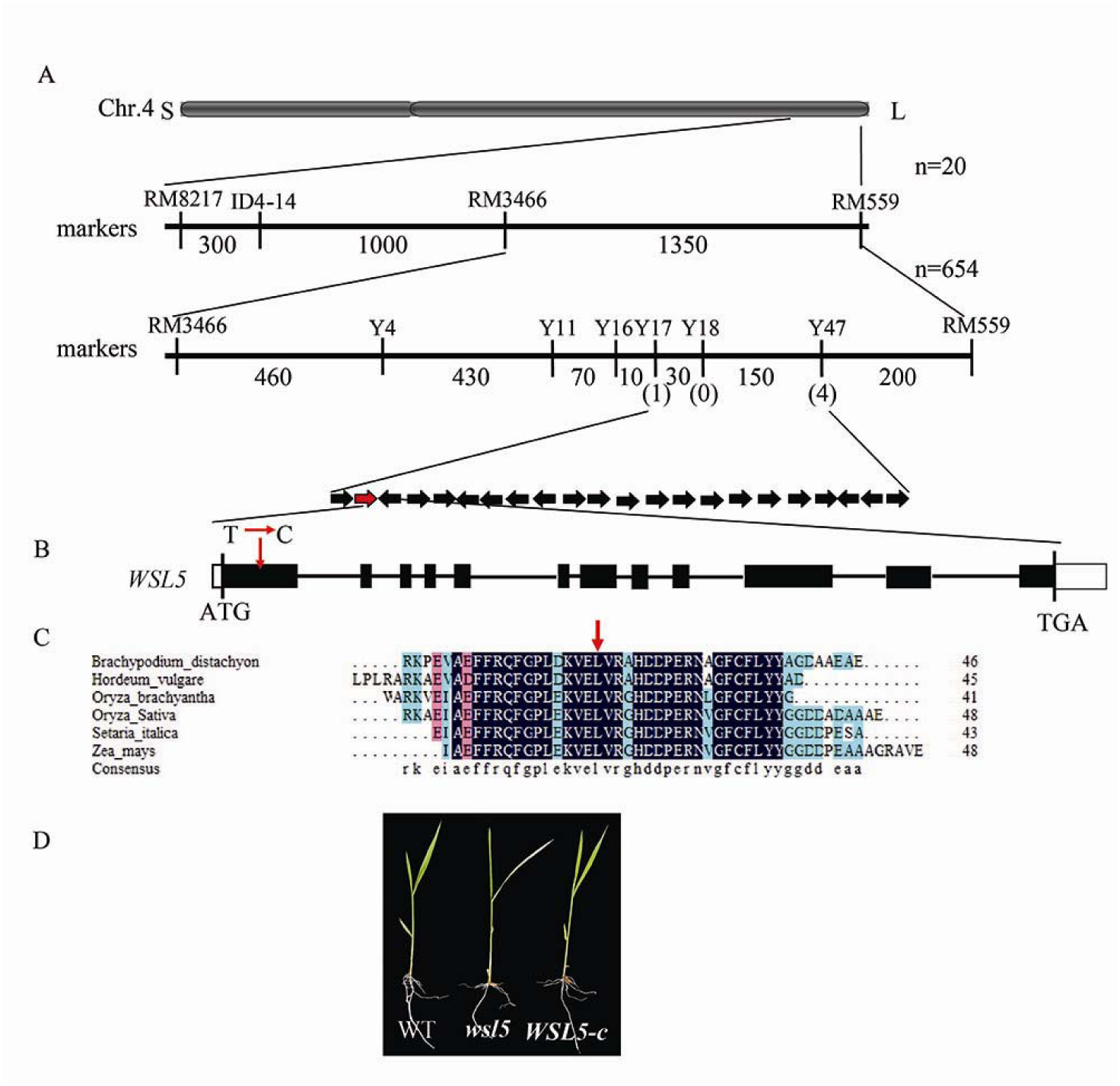
Map-based cloning of the *WSL5* allele. (A) The *WSL5* locus was mapped to a 180 kb region between InDel markers Y17 and Y47, on chromosome 4L. Black arrows represent 22 putative genes in this region; candidate gene *WSL5* (*Os04g0684500*) is shown by a red arrow. (B) ATG and TGA represent the start and stop codons, respectively. Black boxes indicate the exons, and the white boxes indicate the 3′- and 5′- UTR. A SNP in the first exon in *WSL5* causes a leucine to proline amino acid substitution. (C) Alignment of amino acid sequences with highest identity to the WSL5 protein. Red arrow indicates amino acid change. (D) Complementation of *wsl5* by transformation.

To confirm that mutation of *WSL5* was responsible for the mutant phenotype, the *WSL5* coding region driven by the *UBQ* promoter was transformed into calli derived from *wsl5* seeds. Twenty eight of 45 transgenic lines resistant to hygromycin and harboring the transgene displayed the wild-type phenotype (Fig. 3D). These results confirmed that *Os04g0684500* was the *WSL5* gene.

### WSL5 encodes PPR protein

Sequence analysis showed that WSL5 comprised 12 exons and 11 introns. A single base substitution in the *wsl5* mutant was located in the first exon (Fig.3B). A database search with Pfam (http://pfam.xfam.org/search) revealed that WSL5 contained an RNA recognition motif at its N terminus and 15 PPR motifs at the C terminus thus belonging to the P family. The substituted amino acid (Leu) was highly conserved among the RNA recognition motif (Fig. 3C), suggesting an obligate role of this site for functional integrity of WSL5 protein. WSL5 shared a high degree of sequence similarity with maize PPR4 (84% identity) and *Arabidopsis thaliana* At5g04810 (59% identity) (Supplementary Fig. S3). Together, these results indicated that *WSL5* encodes a novel PPR protein.

#### Expression pattern and subcellular localization of WSL5

Using Rice eFP Browser (http://bar.utoronto.ca/efprice/cgi-bin/efpWeb.cgi) we found that *WSL5* was expressed in all tissues, especially in young leaves. To verify these data, we examined the expression levels of *WSL5* in different organs of WT by RT-PCR (Fig. 4A, B). The *WSL5* transcript was preferentially expressed in young leaves (Fig. 4B), suggesting that *WSL5* had an important role in chloroplast development in young seedlings. The *WSL5* transcripts were more abundant in plants grown at 20°C than at 30°C, indicating that *WSL5* was induced by low temperatures. Thus plants might express *WSL5* abundantly to regulate chloroplast development under cold stress (Fig. 4C).

**Fig. 4.**
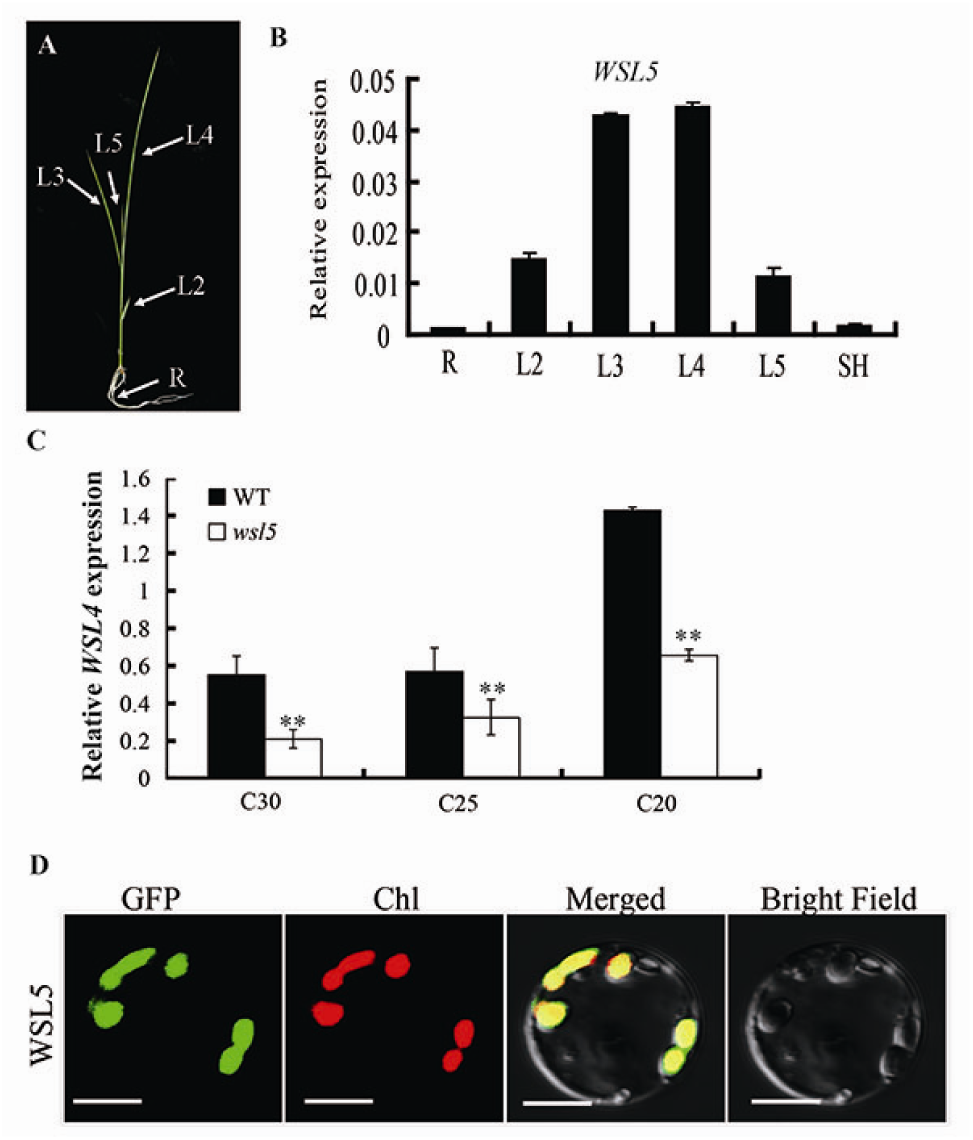
Expression pattern analysis and subcellular localization of WSL5. (A) Schematic of a rice seedling with fully expanded fourth leaf. (B) qRT-PCR analysis *WSL5* expression in roots, stems, L2, L3, L4, L5 and sheaths of wild type. (C) qRT-PCR analyses of *WSL5* transcript in WT and *wsl5* mutant seedlings grown in a growth chamber with a 12 h photoperiod at 30, 25, and 20°C. (D) Localization of WSL5 protein in rice protoplasts. Green fluorescence shows GFP, red fluorescence shows chlorophyll, orange indicates the two types of florescence merged. Error bars represent the SD from three independent experiments. (Student’s t-test, \*\**P* < 0.01).

To examine the actual subcellular localization of WSL5, a CaMV35S-driven construct with a WSL5-GFP fusion protein was generated using the pAN580 vector and transiently expressed in rice protoplasts. Green fluorescent signals of WSL5-GFP co-localized with the autofluorescent signals of chlorophyll (Fig. 4D), suggesting that WSL5 localized to the chloroplasts. These results, together with chloroplast localization and the observed *wsl5* phenotypes, supported the notion that WSL5 plays an important role in regulating chloroplast development in rice seedlings.

#### Expression of photosynthesis related-genes is down-regulated in wsl5

RNA-seq was performed to analyze the effect of the *wsl5* mutation on gene expression. A total of 42 million clean reads were obtained from wild type and *wsl5*. Compared to wild type, there were 1,699 up-regulated genes and 1,999 down-regulated genes in *wsl5* (Fig. 5A-C and Supplementary Data S1). We randomly selected 5 down-regulated and 5 up-regulated genes to verify the results of RNA-seq. The qRT-PCR results were consistent with those from RNA-seq (Fig. 5D). Go and KEGG enrichment analysis indicated that genes encoding photosynthesis, light reaction, PSI and PSII, chloroplast thylakoid, ATP synthase, and carbon fixation had reduced expression in *wsl5* (Supplementary Fig. S5 and S6). Also, some chlorophyll synthesis genes, including *HEMA, YGL8, PORA, CHLH, CRD1*, were significantly reduced, which was verified using real-time PCR (Supplementary Fig. S7).

**Fig. 5.**
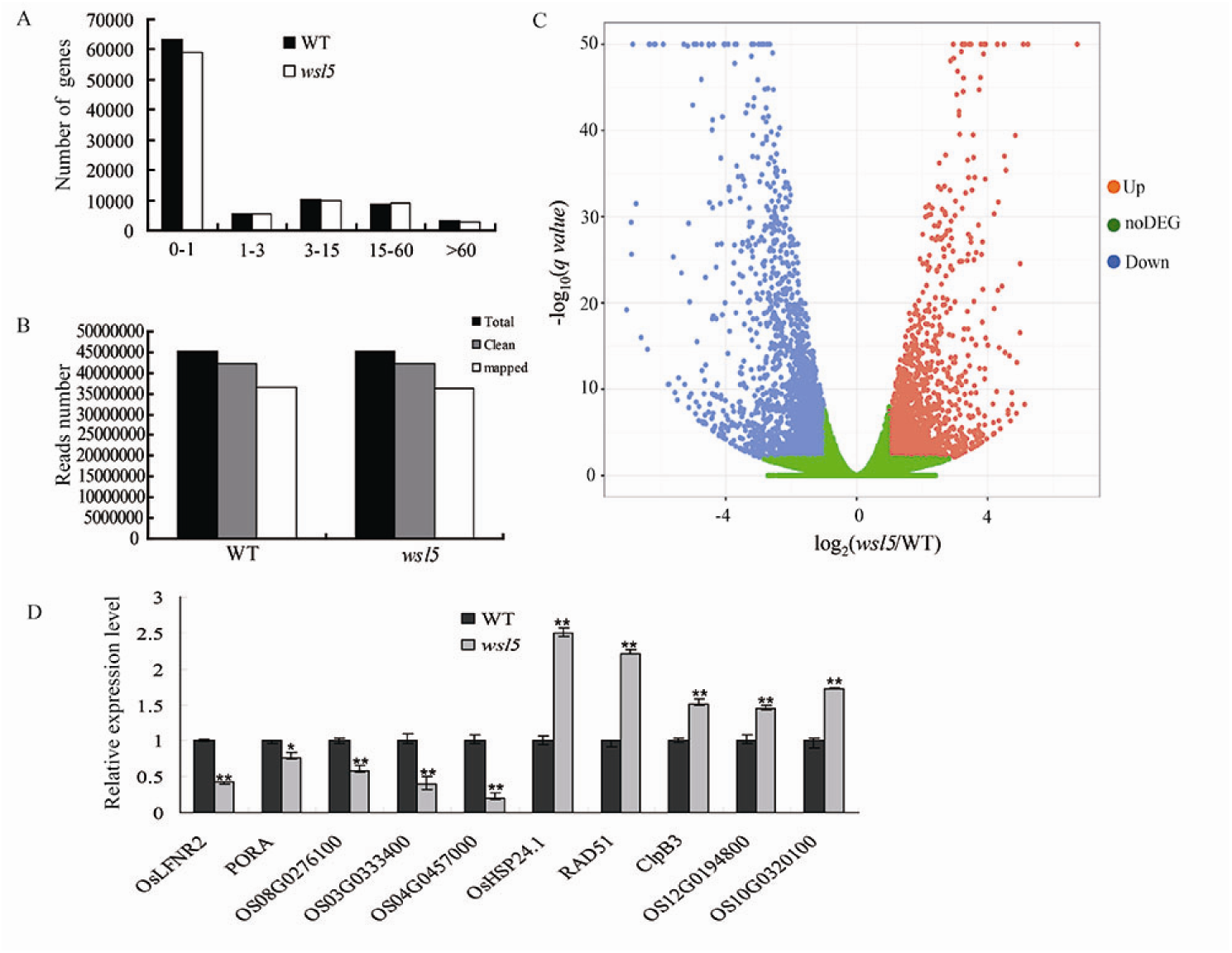
RNA-seq analysis of wild-type and *wsl5* seedlings. mRNA was enriched from total RNA isolated from 10-d-old (third leaf) seedlings of wild type and *wsl5* using oligo-(dT). cDNA was synthesized using random hexamer primers and reverse-transcribed using random hexamer primers. The library was then constructed and sequenced using an Illumina HiSEquation 2000. (A) Frequencies of detected genes sorted according to expression levels. (B) Read numbers of wild-type and *wsl5* sequences. (C) Volcano plot showing the overall alterations in gene expression in wild type and *wsl5*. (D) qRT-PCR analysis of genes differentially expressed in RNA-seq. Five up-regulated and 5 down-regulated genes were tested. Error bars represent SD from three independent experiments. (Student’s t-test, \**P* < 0.05, \*\**P* < 0.01).

### wsl5 mutants have global defects in plastid gene expression

To investigate whether the *WSL5* mutation affects transcription by PEP and NEP, we examined transcript abundance of various plastid genes in the *wsl5* mutant by RNA-seq. The expression levels of many plastidic genes differed between *wsl5* and wild type (Fig. 6). Compared with the wild type, the expressions of the plastid genes that are transcribed by PEP, including *psbA, psbB, psbD, petB, ndhA*, and *rbcL*, were strongly reduced in *wsl5* mutant. In addition, transcript levels of the plastid genes, including the *ribosomal protein L32* (*rpl32*)*, rpl14, rps2, rps4*, and *rpoA*, which are transcribed by NEP, were increased or unchanged in the mutant (Fig. 6). These results indicated that *WSL5* was required for optimal expression of plastid genes in rice seedlings.

**Fig. 6.**
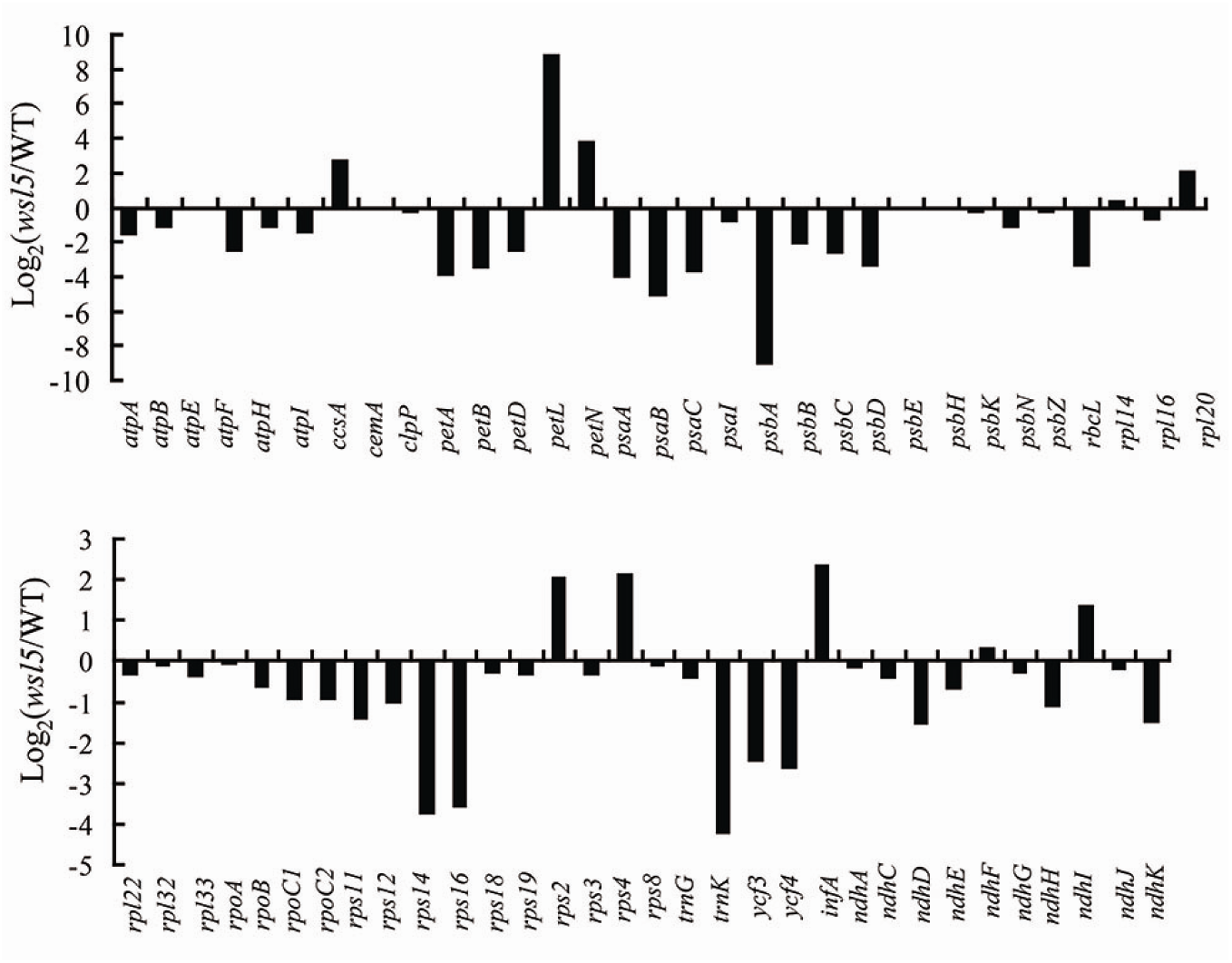
Differential expression of plastid-encoded genes in wild type and *wsl5*. mRNA enriched from total RNA isolated from 10-d-old seedlings of wild type and *wsl5* was fragmented and reverse-transcribed using random hexamer primers. The library was then constructed and sequenced using an Illumina HiSEquation 2000. The graph shows the log_2_ ratio of transcript levels in *wsl5* mutant compared with wild type.

### Analysis of transcripts and proteins of genes associated with chloroplast biogenesis in wsl5

Since WSL5 was located in chloroplasts, we tested the accumulation of chloroplast proteins in *wsl5* and wild type using western-blot analysis under the C20 and C30 conditions. Under the C20 condition, protein levels in the large subunit of Rubisco (RbcL) and Rubisco activase (RCA) were much lower in *wsl5* (Fig. 7A). Other plastidic proteins including NADH dehydrogenase subunit 4, A1 of PSI, D1 of PSII, alpha subunit of RNA polymerase were tested. The results showed that the levels of plastid-encoded proteins were significantly decreased in *wsl5* (Fig. 7A). qRT-PCR results suggested the expression levels of class I genes *RbcL, psbA, psaA* were strikingly reduced, whereas expression of class III genes *rpoA* and *rpoC1*, and class II gene *AtpB*, was unchanged (Fig. 7B). When grown in C30 conditions, transcripts and proteins of all genes in the mutant and WT showed very slight differences in expression pattern (Fig. 7C-D). These results indicated that *WSL5* was required for protecting PEP activity under cold stress.

**Fig. 7.**
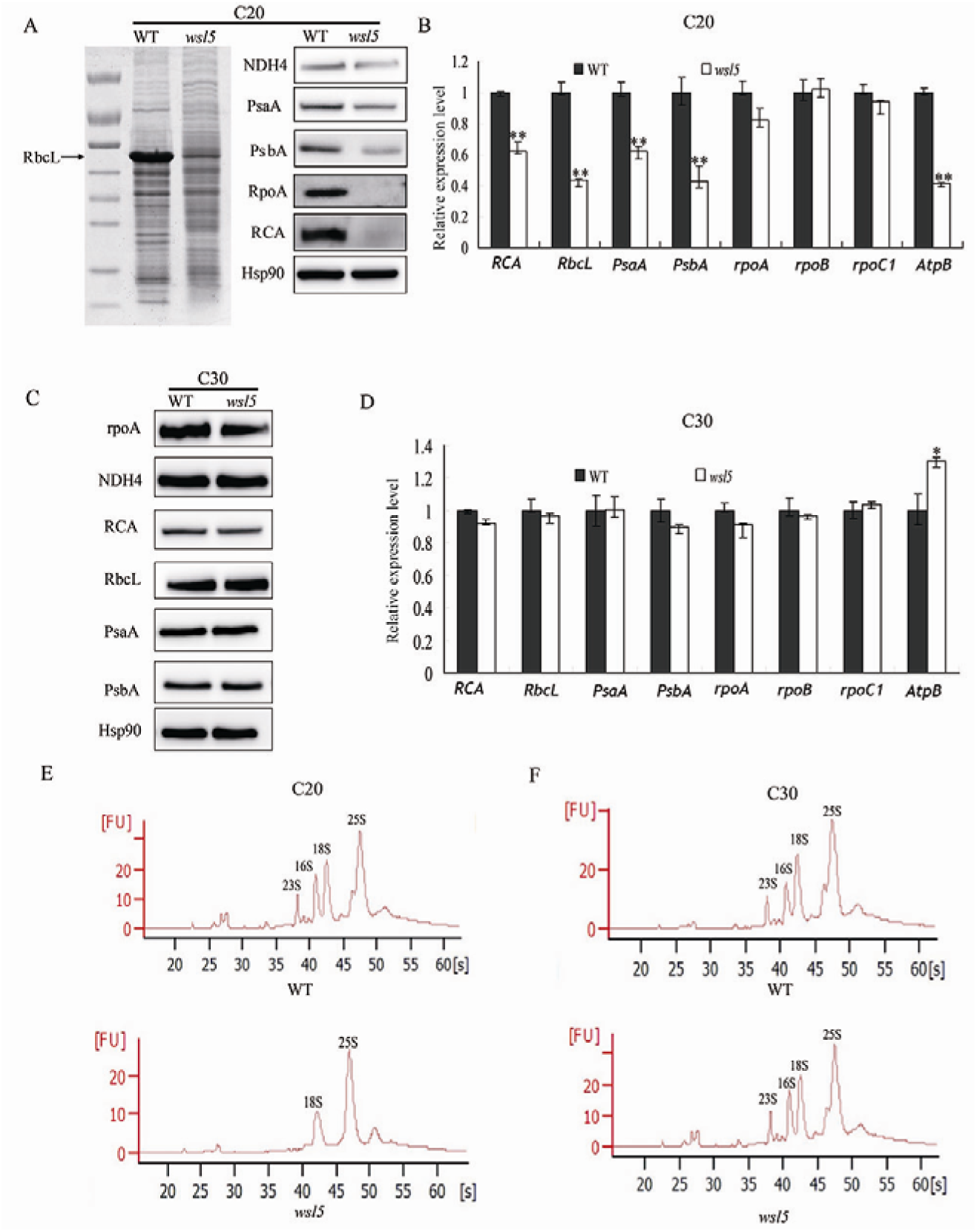
Analysis of accumulation of transcripts and proteins of representative genes associated with chloroplast biogenesis in WT and *wsl5* seedlings. (A, C) Western blot analysis of chloroplast proteins and RCA in wild-type and *wsl5* mutant seedlings at the third-leaf stage at C20 (A) and C30 (C). Hsp90 was used as an internal control. (B, D) qRT-PCR analysis of relative expression levels of plastidic encoding genes in wild type and *wsl5* at the third-leaf stage under (B) C20 or (D) C30. Error bars represent SD from three independent experiments. (E-F) rRNA analysis using an Agilent 2100 bioanalyzer. RNA was isolated from 10-d-old wild-type seedlings and *wsl5* seedlings grown in C30 and C20. (Student’s t-test, \*\**P* < 0.01).

The chloroplast ribosome consists of a 50S large subunit and a 30S small subunit. Both subunits are comprised of rRNAs (23S, 16S, 5S, and 4.5S) and ribosomal proteins. We analyzed the composition and content of rRNAs using an Agilent 2100 bioanalyzer under C20 and C30 conditions. rRNA, including the 23S and 16S rRNAs, were decreased in *wsl5* seedlings under cold stress, but there was no difference under C30 conditions (Fig. 7E, F). These results clearly indicated severe defects in plastidic ribosome biogenesis in the *wsl5* mutant seedlings grown in cold conditions.

### The wsl5 mutant is defective in RNA editing and splicing of chloroplast group II introns

PPR proteins are required for RNA editing, splicing, stability, maturation, and translation (Tan *et al*. 2014; Hammani *et al*. 2016). Since WSL5 belongs to the P group, it was likely involved in transcript-processing activities. Firstly, we determined whether loss of *WSL5* function affected editing at 21 identified RNA editing sites in chloroplast RNA (Corneille *et al*. 2000). The results showed that the editing efficiencies of *rpl2* at C1 and *atpA* at C1148 were significantly decreased in *wsl5* mutant compared to WT (Supplementary Fig.S8), whereas the other 10 genes and corresponding 19 editing sites were normally edited in *wsl5* mutant. We then analyzed the editing efficiencies of *rpl2* at C1 and *atpA* at C1148 in complemented transgenic plants. As expected, the editing efficiencies of *rpl2* at C1 and *atpA* at C1148 were markedly improved in complemented plants (Supplementary Fig.S8). These data supported the contention that the mutation in WSL5 affected the editing efficiency of *rpl2* at C1 and *atpA* at C1148.

In *Arabidopsis thaliana*, multiple organellar RNA editing factor (MORF) proteins have been implicated in RNA editing and provide the link between PPR proteins and the proteins contributing the enzymatic activity (Takenaka *et al*. 2012). Based on *Arabidopsis thaliana* MORF protein families (Zehrmann *et al*. 2015), we examined the potential interactions between rice MORF proteins and WSL5 by yeast two-hybrid analysis. The results showed that Os09g33480 and Os09g04670, both belonging to the *Arabidopsis thaliana* MORF8 branch (Zhang *et al*. 2017), strongly interacted with WSL5 protein in yeast (Fig.8). In contrast, Os04g51280, Os06g02600 and Os08g04450 did not interact with WSL5 (Fig.8). These results suggested that *WSL5* may participate in RNA editing by interacting with *OsMORF8s*.

**Fig. 8.**
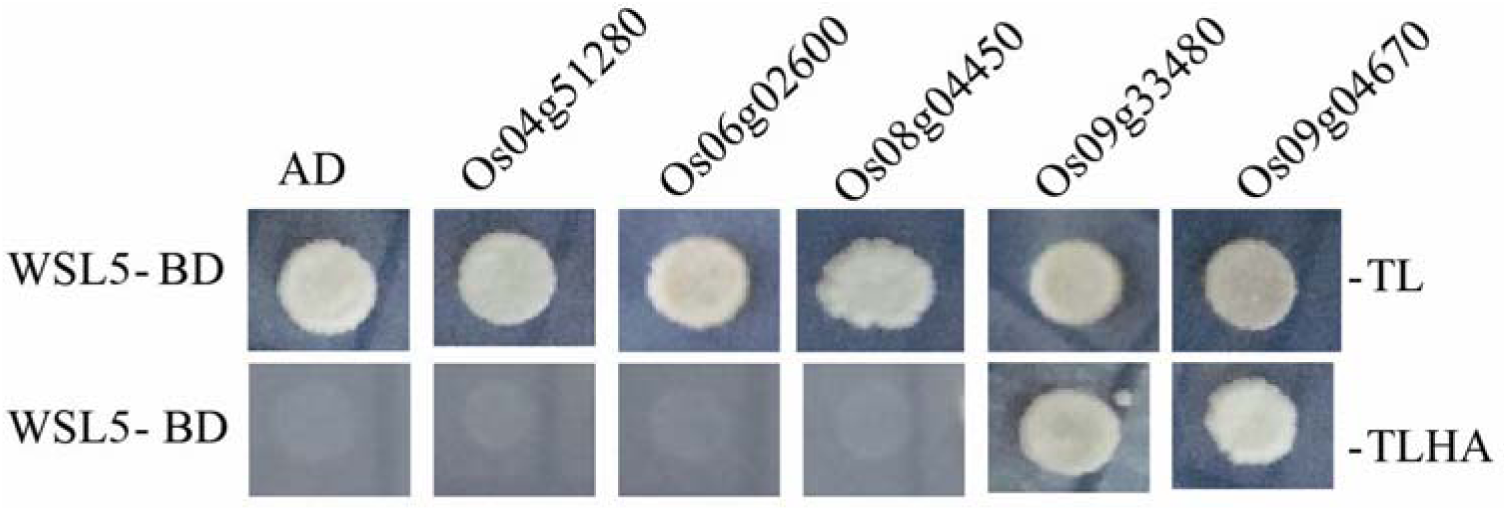
Yeast two-hybrid assay of WSL5 and MORF families. WSL5 was fused to the pGBKT7 vector (WSL5-BD). MORF protein was fused to the pGADT7 vector. LT, control medium (SD–Leu/–Trp); LTHA, selective medium (SD–Leu/–Trp/–His/–Ade). Empty pGBKT7 and pGAD-T7 vectors served as negative controls.

We tested whether WSL5 is involved in RNA splicing of chloroplast genes. The rice chloroplast genome contains 18 introns (17 group II introns and one group I intron) (Kaminaka *et al*. 1999). We amplified all chloroplast genes with at least one intron by RT-PCR using primers flanking the introns and compared the lengths of the amplified products between WT and *wsl5* mutant. Chloroplast transcripts *rpl2* and *rps12-2* were spliced at very low efficiency in *wsl5* compared to WT (Fig. 9, 10 and Supplementary Fig. S9). To gain insight into the effects of the impaired splicing of *rpl2* and *rps12-2* on post-processing, we performed qRT-PCR to examine the expression of *rpl2* and *rps12* in *wsl5*. The *rpl2* and *rps12* transcript abundances were high in the mutant compared to WT (Fig. 10C, D). Thus, the low splicing efficiency of *rpl2* and *rps12-2* resulted in aberrant transcript accumulation in the *wsl5* mutant.

**Fig. 9.**
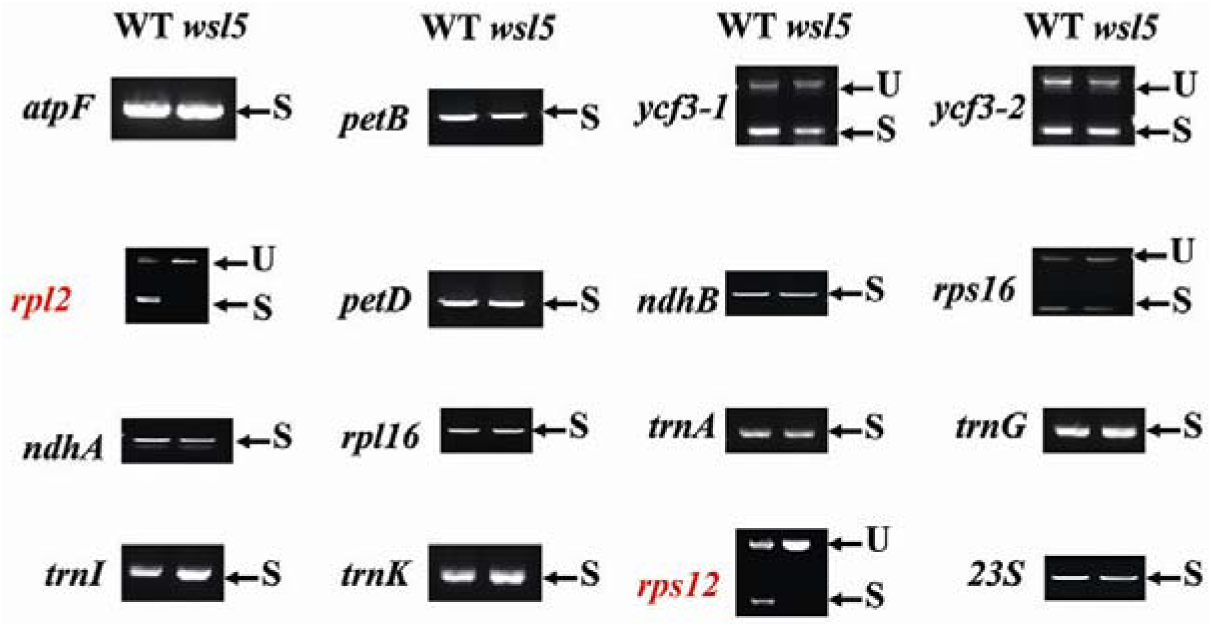
Splicing analyses of chloroplast transcripts in WT and *wsl5* mutant. Gene transcripts are labeled at the left. Spliced (S) and unspliced (U) transcripts are shown at the right. RNA was extracted from WT and *wsl5* mutant seedlings.

**Fig. 10.**
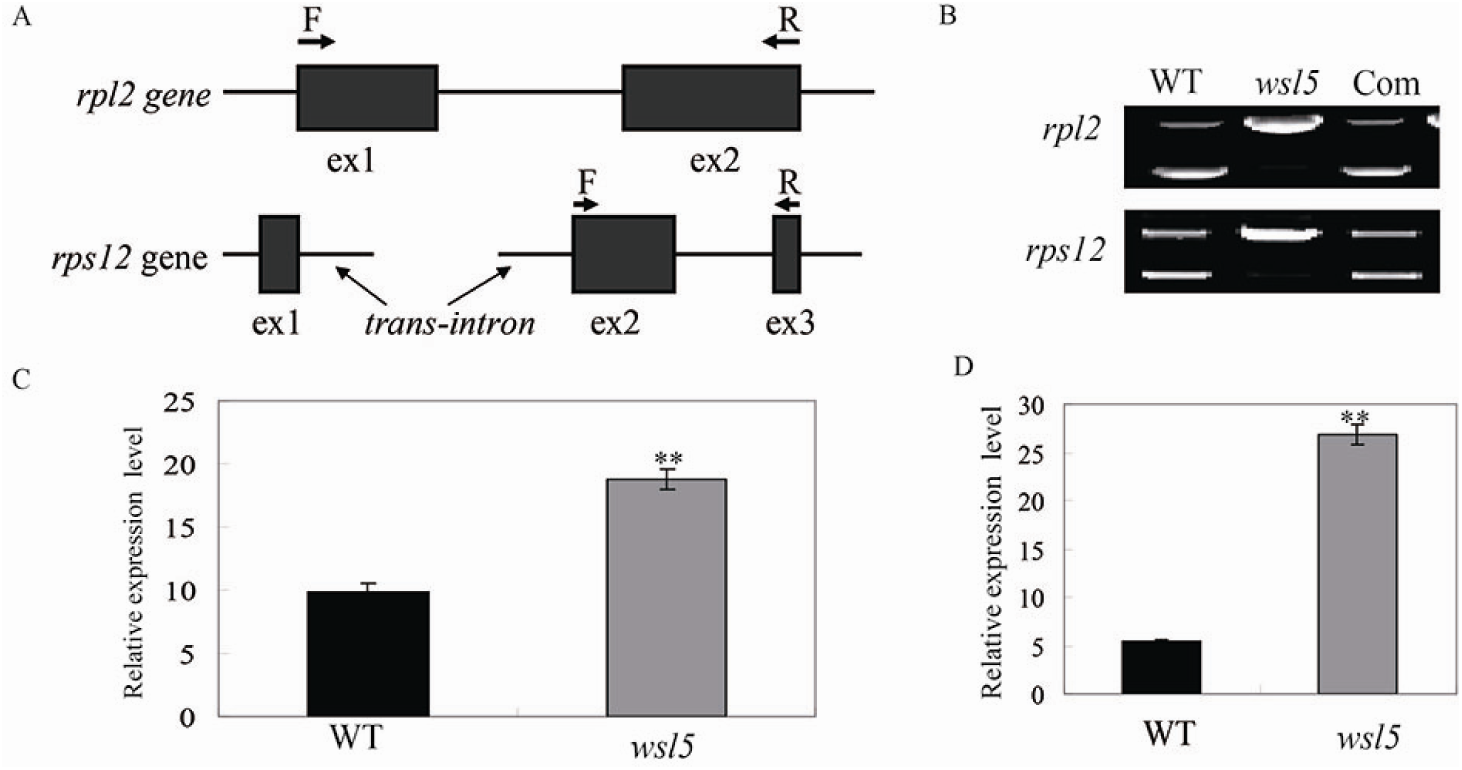
Splicing analyses of two chloroplast group II introns in WT and *wsl5* mutant. (A) Sketch map of *rpl2* and *rps12* transcripts. (B) RT-PCR analyses of *rpl2* and *rps12* transcripts in WT and *wsl5* mutant. (C-D) Quantitative RT-PCR analyses of *rpl2* and *rps12* transcripts in WT and *wsl5* mutant seedlings. Data are means ± SD of three repeats. Student’s t-test: \*\**P* < 0.01.

### Differentially expressed gene analysis in wsl5 and wild type under cold stress and normal conditions

To investigate why phenotypic variation in *wsl5* mutants depends on temperature, we carried out differential gene expression analysis of *wsl5* and WT seedlings grown in growth cabinets at C20 and C30 by RNA-seq. mRNA was purified from total RNA isolated from the third leaves using poly-T oligo-attached magnetic beads; 6,491 overlapping genes were up or down-regulated between the two temperature treatments (Fig. 11 A, B and Supplementary Data S2). Go analysis indicated that genes involved in metabolic processes, oxidation-reduction processes, photosynthesis, light reaction, PSI and PSII, chloroplast thylakoid, ATP synthase, and carbon fixation were strongly reduced in the *wsl5* mutant at C20 (Fig. 11C). These results indicated that the *WSL5* mutation led to change in many physiological processes under cold stress.

**Fig. 11.**
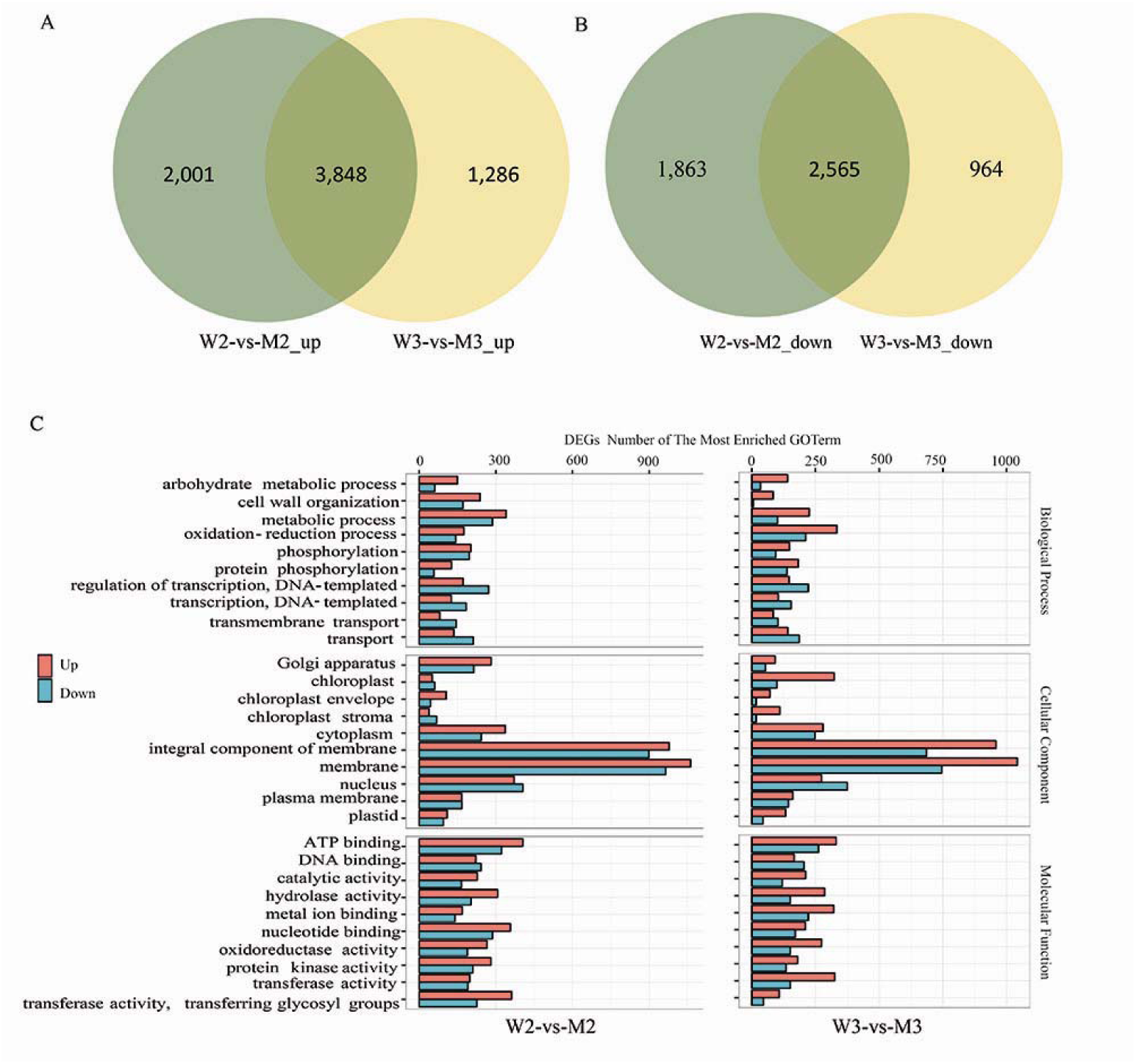
RNA-seq analysis of wild type and *wsl5* under low temperature and normal conditions. (A) Up-regulated differentially expressed genes comparing W2 and M2 and W3 and M3. (B) Down-regulated differentially expressed genes between W2-vs-M2 and W3-vs-M3. (C) Go analysis of genes differentially expressed between W2 and M2 and W3 and M3. W3 and W2 represent wild type plants grown at 30°C and 20°C, respectively. M3 and M2 represent *wsl5* plants grown at 30°C and 20°C, respectively.

## Discussion

### WSL5 encodes a chloroplast-targeted PPR protein that is essential for chloroplast development in juvenile plants under cold stress

PPR genes constitute a large multigene family in higher plants. PPR proteins are essential for plant growth and development and most of them are involved in RNA editing, splicing, and regulation of stability of various organellar transcripts (Barkan and Small, 2014). In contrast to PPRs in *Arabidopsis thaliana*, little is known about the functions of PPRs in rice. Here, we present a molecular characterization of PPR gene *WSL5* in rice. It has an RNA recognition motif and 15 PPR motifs (Supplementary Fig.S3). The WSL5 protein was predicted to contain a chloroplast transit peptide (cTP) in its N-terminal region, suggesting that the protein is one of the PPRs targeted to chloroplasts, and this was confirmed by subcellular localization experiments (Fig. 4D). The disruption of WSL5 protein under natural conditions led to abnormal chloroplasts and caused a variegated phenotype that affected both the chlorophyll content and the chloroplast ultrastructure up to the fourth leaf growth stage, whereas the *wsl5* mutant was albinic under cold stress (Fig. 2 and Supplementary Fig. S2). This finding suggests that the function of WSL5 is essential for early chloroplast development under cold stress in rice. This conclusion is further supported by the results of expression analysis. *WSL5* was highly expressed in leaf section L3 and L4 at the seedling stage. A high level of WSL5 was noted under low temperature. Sequence alignment of homologous proteins using *Arabidopsis thaliana*, maize and rice showed that the mutant site in *wsl5* is conserved within the RRM motif.

### WSL5 is involved in splicing of plastid genes and in ribosome biosynthesis

A large group of nuclear-encoded PPR proteins involved in RNA editing, splicing, stability, maturation, and translation is required chloroplast development (Tan *et al*. 2014; Wang *et al*. 2017). To date, six PPR proteins have been reported to be involved in RNA splicing of group II introns in chloroplasts. Among them, the maize PPR4 protein acts as an *rps12* trans-splicing factor (Schmitz-Linneweber *et al*. 2006). *Arabidopsis thaliana* PPR protein OTP51 functions as a plastid *ycf3-2* intron cis-splicing factor and OTP70 has been implicated in splicing of the plastid transcript *rpoC1* (de Longevialle *et al*. 2008; Chateigner-Boutin *et al*. 2011). In this study, the *wsl5* mutant caused defects in splicing of *rpl2* and *rps12* (Figs. 9 and 10), implying that WSL5 probably controls chloroplast RNA intron splicing during early leaf development in rice. This finding indicates that disruption of *rpl2* or *rps12*, either alone or in combination, may be responsible for the mutant phenotype.

Defective *rps12* and *rpl2* splicing could account for the white-stripe leaf phenotype and plastid ribosome deficiency in *wsl5* mutant (Figs. 9 and 10). We analyzed the contents of rRNAs and ribosomal proteins; 23S and 16S rRNAs were decreased in *wsl5* mutant under cold stress (Fig. 7E-F). The lack of mature *rps12* and *rpl2* mRNA in *wsl5* mutants may severely affect ribosome functions in plastids. Thus, the ribosome assembly defect in *wsl5* may also contribute to the *wsl5* phenotype.

### Possible mechanism of WSL5 regulating chloroplast development under cold stress and normal conditions

To study the molecular mechanism of *WSL5* in regulating chloroplast development under different temperature conditions we compared gene expression patterns in *wsl5* mutant and wild type by RNA-seq analysis. Our findings showed that under cold stress *WSL5* regulates expression of genes, involved in carbohydrate metabolic processes, oxidation-reduction processes, photosynthesis, biosynthesis of secondary metabolites, chlorophyll biosynthesis process, and chloroplast development (Fig. 11 and Supplementary Data S2). Plastid thioredoxins are important for maintaining plastid oxidation-reduction balance (Bohrer *et al*. 2012). Many genes involved in regulating plastid oxidation-reduction balance are changed under the C20 and C30 conditions, such as *OsTRXm, OsTRXz* (Supplementary Fig. S10). *OsTRXm* is involved in regulation of activity of a target peroxiredoxin (Prx) through reduction of Cystic disulfide bridges (Chi *et al*. 2008). OsTRXz interacts with TSV to protect chloroplast development under cold stress (Sun *et al*. 2017). The large and small subunits of ribonucleotide reductase (RNR), V3 and St1 regulate the rate of deoxyribonucleotide production for DNA synthesis and repair (Yoo *et al*. 2009). *V3* and *St1* are repressed under constant 20□ conditions in *wsl5*, indicating that mutation in *WSL5* leads to defects in DNA synthesis and repair in juvenile plants at low temperatures (Supplementary Fig. S10). The expression levels of fatty acid metabolism genes *OsFAH1, OsFAH2*, and *OsFAD7*, and plastid starch metabolism genes *AGPS2b* and *PHO1*, were all dramatically changed in *wsl5* compared with wild-type at low temperatures (Supplementary Fig. S10). These results indicate that *WSL5* is essential for chloroplast development under cold stress.

In conclusion, *WSL5* plays an important role in expression of plastid genes and biogenesis of plastid ribosomes, and is essential for chloroplast development in rice seedlings under cold stress by coordinated transcription and translation of chloroplast-associated genes. Identification of this new PPR protein will help to elucidate the molecular mechanisms of plastid development and ribosome biogenesis, and shed light on understanding chloroplast development in juvenile plants grown under cold stress.

## Supplementary data

Additional supplementary data may be found online for this article:

**Data S1** Genes differentially expressed in wild type and *wsl5*.

**Data S2** Genes differentially expressed in wild type and *wsl5* under different temperature conditions.

**Fig. S1.** Comparison of pigment contents from the second (L2), third (L3), fourth (L4) and fifth (L5) leaves of five-leaf-stage plants between WT and *wsl5* mutant.

**Fig. S2.** Transmission electron microscopy images of cells from WT and *wsl5* mutants grown under different temperature conditions.

**Fig. S3.** Alignment of *WSL5* orthologs in maize and *Arabidopsis*.

**Fig. S4.** *WSL5* was expressed in all tissues, especially during leaf development according to Rice eFP Browser.

**Fig. S5.** GO analysis of genes differentially expressed between wild type and *wsl5*.

**Fig. S6.** Pathway analysis of genes differentially expressed between wild type and *wsl5*.

**Fig. S7.** Expression levels of chlorophyll synthesis genes in wild type and *wsl5*.

**Fig. S8.** Editing efficiencies of *rpl2* and *atpA* genes in WT and the *wsl5* mutant.

**Fig. S9.** Quantitative RT-PCR analyses of *rpl2*, and *rps12* transcripts in WT and the *wsl5* mutant.

**Fig. S10.** qRT-PCR analysis of genes differently expressed in RNA-seq.

**Table S1.** Comparison of agronomic traits between WT and *wsl5* under field conditions.

**Table S2.** Primers used in this study.

## Acknowledgements

This research was supported by Key Laboratory of Biology, Genetics and Breeding of Japonica Rice in Mid-lower Yangtze River, Ministry of Agriculture, P.R. China, and Jiangsu Collaborative Innovation Center for Modern Crop Production, and grants from The National Key Research and Development Program of China (2016YFD0101801, 2016YFD0100101-08), Jiangsu Science and Technology Development Program (BE2017368), Agricultural Science and Technology Innovation Fund project of Jiangsu Province(CX(16)1029) and Key Program of Science and Technology of Anhui Province

## References

Arnon DI. 1949. Copper enzymes in isolated chloroplasts. Polyphenoloxidase in *Beta vulgaris*. Plant Physiology 24, 1–15.

Barkan A, Small I. 2014. Pentatricopeptide repeat proteins in plants. Annual Review of Plant Biology 65, 415–442.

Bohrer AS, Massot V, Innocenti G, Reichheld JP, Issakidis-Bourguet E, Vanacker H. 2012. New insights into the reduction systems of plastidial thioredoxins point out the unique properties of thioredoxin z from Arabidopsis. Journal of Experimental Botany 63, 6315–6323.

Chateigner-Boutin AL, des Francs-Small CC, Delannoy E, Kahlau S, Tanz SK, de Longevialle AF, Fujii S, Small I. 2011. OTP70 is a pentatricopeptide repeat protein of the E subgroup involved in splicing of the plastid transcript *rpoC1*. Plant Journal 65, 532–542.

Chen S, Tao L, Zeng L, Vega-Sanchez ME, Umemura K, Wang GL. 2006. A highly efficient transient protoplast system for analyzing defence gene expression and protein-protein interactions in rice. Molecular Plant Pathology 7, 417–427.

Chi YH, Moon JC, Park H, et al. 2008. Abnormal chloroplast development and growth inhibition in rice thioredoxin m knock-down plants. Plant Physiology 148, 808–817.

Corneille S, Lutz K, Maliga P. 2000. Conservation of RNA editing between rice and maize plastids: are most editing events dispensable? Molecular Genetics and Genomics 264, 419–424.

de Longevialle AF, Hendrickson L, Taylor NL, Delannoy E, Lurin C, Badger M, Millar AH, Small I. 2008. The pentatricopeptide repeat gene *OTP51* with two LAGLIDADG motifs is required for the cis-splicing of plastid *ycf3* intron 2 in *Arabidopsis thaliana*. Plant Journal 56, 157–168.

Gong X, Su Q, Lin D, Jiang Q, Xu J, Zhang J, Teng S, Dong Y. 2014. The rice *OsV4* encoding a novel pentatricopeptide repeat protein is required for chloroplast development during the early leaf stage under cold stress. Journal of Integrative Plant Biology 56, 400–410.

Hammani K, Takenaka M, Miranda R, Barkan A. 2016. A PPR protein in the PLS subfamily stabilizes the 5’-end of processed *rpl16* mRNAs in maize chloroplasts. Nucleic Acids Research 44, 4278–4288.

Hedtke B, Borner T, Weihe A. 1997. Mitochondrial and chloroplast phage-type RNA polymerases in *Arabidopsis*. Science 277, 809–811.

Ichinose M, Tasaki E, Sugita C, Sugita M. 2012. A PPR-DYW protein is required for splicing of a group II intron of *cox1* pre-mRNA in *Physcomitrella patens*. Plant Journal 70, 271–278.

Jarvis P, López-Juez E. 2013. Biogenesis and homeostasis of chloroplasts and other plastids. Nature Review Molecular Cell Biology 14, 787–802.

Jiang Q, Mei J, Gong XD, Xu JL, Zhang JH, Teng S, Lin DZ, Dong YJ. 2014. Importance of the rice *TCD9* encoding α subunit of chaperonin protein 60 (Cpn60α) for the chloroplast development during the early leaf stage. Plant Science 215–216, 172–179.

Kaminaka H, Morita S, Tokumoto M, Yokoyama H, Masumura T, Tanaka K. 1999. Molecular cloning and characterization of a cDNA for an iron-superoxide dismutase in rice (*Oryza sativa* L.). Bioscience. Biotechnology. Biochemistry 63, 302–308.

Kanehisa M, Araki M, Goto S, et al. 2008. KEGG for linking genomes to life and the environment. Nucleic Acids Research 36, 480–484.

Kusumi K, Iba K. 2014. Establishment of the chloroplast genetic system in rice during early leaf development and at low temperatures. Frontiers in Plant Science 5, 386.

Liere K, Weihe A, Börner T. 2011. The transcription machineries of plant mitochondria and chloroplasts: composition, function, and regulation. Journal of Plant Physiology 68, 1345–1360.

Lin D, Gong X, Jiang Q, Zheng K, Zhou H, Xu J, Teng S, Dong Y. 2015. The rice *ALS3* encoding a novel pentatricopeptide repeat protein is required for chloroplast development and seedling growth. Rice 8, 17.

Liu F, Xu W, Song Q, Tan L, Liu J, Zhu Z, Fu Y, Su Z, Sun C. 2013. Microarray assisted fine-mapping of quantitative trait loci for cold tolerance in rice. Molecular Plant 6, 757–767.

O’Toole N, Hattori M, Andres C, Iida K, Lurin C, Schmitz-Linneweber C, Sugita M, Small I. 2008. On the expansion of the pentatricopeptide repeat gene family in plants. Molecular Biology and Evolution 25, 1120–1128.

Schmitz-Linnerweber C, Williams-Carrier Re, Kroeger TS, Vichas A, Barkan A. 2006. A pentatricopeptide repeat protein facilitates the trans-splicing of the maize chloroplast rps12 pre-mRNA. Plant Cell 18, 2650–2663.

Shikanai T, Fujii S. 2013. Function of PPR proteins in plastid gene expression. RNA Biology 10, 1446–1456.

Stern DB, Goldschmidt-Clermont M, Hanson MR. 2010. Chloroplast RNA metabolism. Annual Review of Plant Biology 61, 125–155.

Su N, Hu ML, Wu DX, et al. 2012. Disruption of a rice pentatricopeptide repeat protein causes a seedling-specific albino phenotype and its utilization to enhance seed purity in hybrid rice production. Plant Physiology 159, 227–238.

Sun J, Zheng T, Yu J, et al. 2017. TSV, a putative plastidic oxidoreductase, protects rice chloroplasts from cold stress during development by interacting with plastidic thioredoxin Z. New Phytologist 215, 240–255.

Takenaka M, Brenniche A. 2007. RNA editing in plant mitochondria: assays and biochemical approaches. Methodsin Enzymology 424, 439–458.

Takenaka M, Zehrmann A, Verbitskiy D, Hartel B, Brennicke A. 2013. RNA editing in plants and its evolution. Annual Review of Genetics 47, 335–352.

Takenaka M, Zehrmann A, Verbitskiy D, Kugelmann M, Hartel B, Brennicke A. 2012. Multiple organellar RNA editing factor (MORF) family proteins are required for RNA editing in mitochondria and plastids of plants. Proceedings of the National Academy of Sciences of the United States of America 109, 5104–5109.

Tan J, Tan A, Wu F, et al. 2014. A novel chloroplast-localized pentatricopeptide repeat protein involved in splicing affects chloroplast development and abiotic stress response in rice. Molecular Plant 7, 1329–1249.

Tillich M, Krause K. 2010. The ins and outs of editing and splicing of plastid RNAs: lessons from parasitic plants. Nature Biotechnology 27, 256–266.

Wang D, Liu H, Zhai G, Wang L, Shao J, Tao Y. 2016. *OspTAC2* encodes a pentatricopeptide repeat protein and regulates rice chloroplast development. Journal of Genetics and Genomics 43, 601–608.

Wang Y, Ren Y, Zhou K, et al. 2017. *WHITE STRIPE LEAF4* encodes a novel P-type PPR Protein required for chloroplast biogenesis during early leaf development. Frontiers in Plant Science doi:10.3389/fpls.2017.01116.

Wang Y, Wang C, Zheng M, et al. 2016. WHITE PANICLE1, a val-tRNA synthetase regulating chloroplast ribosome biogenesis in rice, is essential for early chloroplast development. Plant Physiology 170, 2110–2123.

Yoo SC, Cho SH, Sugimoto H, Li J, Kusumi K, Koh HJ, Iba K, Paek NC. 2009. Rice *Virescent3* and *Stripe1* encoding the large and small subunits of ribonucleotide reductase are required for chloroplast biogenesis during early leaf development. Plant Physiology 50, 388–401.

Young MD, Wakefeld MJ, Smyth GK, Oshlack A. 2010. Gene ontology analysis for RNA-seq: accounting for selection bias. Genome Biology 11, R14.

Yu Q.B, Huang C, Yang ZN. 2014. Nuclear-encoded factors associated with the chloroplast transcription machinery of higher plants. Frontiers in Plant Science 5, 316.

Yu QB, Jiang Y, Chong K, Yang ZN. 2009. AtECB2, a pentatricopeptide repeat protein, is required for chloroplast transcript accD RNA editing and early chloroplast biogenesis in *Arabidopsis thaliana*. Plant Journal 59, 1011–1023.

Zehrmann A, Härtel B, Glass F, Bayer-Császár E, Obata T, Meyer E, Brennicke A, Takenak M. 2015. Selective homo- and heteromer interactions between the multiple organellar RNA editing factor (MORF) proteins in *Arabidopsis thaliana*. Journal of Biological Chemistry 290, 6445–6456.

Zhang Z, Cui X, Wang Y, Wu J, Gu X, Lu T. 2017. The RNA editing factor *WSP1* is essential for chloroplast development in rice. Molecular Plant 10, 86–89.

Zhao J, Zhang S, Dong J, Yang T, Mao X, Liu Q, Wang, X., Liu B. 2017. A novel functional gene associated with cold tolerance at the seedling stage in rice. Plant Biotechnology Journal 15, 1141–1148.

